# A genome-wide association study identifies markers and candidate genes affecting tolerance to the wheat pathogen *Zymoseptoria tritici*

**DOI:** 10.1101/2024.08.08.607179

**Authors:** Alexey Mikaberidze, Bruce A. McDonald, Lukas Kronenberg

**Affiliations:** School of Agriculture, Policy and Development; University of Reading, Reading, RG6 6EU, United Kingdom; Plant Pathology, Institute of Integrative Biology, ETH Zurich, Zurich, 8092, Switzerland; Crop Genetics, The John Innes Centre, Norwich, NR4 7UH, United Kingdom

**Keywords:** leaf tolerance, trade-off, GWAS, disease resistance, marker-trait association, candidate genes, *Zymoseptoria tritici*, septoria tritici blotch, plant defense

## Abstract

Plants defend themselves against pathogens using either resistance, measured as the host’s ability to limit pathogen multiplication, or tolerance, measured as the host’s ability to reduce the negative effects of infection. Tolerance is a promising trait for crop breeding, but its genetic basis has rarely been studied and remains poorly understood. Here, we reveal the genetic basis of leaf tolerance to the fungal pathogen *Zymoseptoria tritici* that causes the globally important septoria tritici blotch disease on wheat. Leaf tolerance to *Z. tritici* is a quantitative trait that was recently discovered in wheat by using automated image analyses that quantified the symptomatic leaf area and counted the number of pycnidia found on the same leaf. A genome-wide association study identified four chromosome intervals associated with tolerance and a separate chromosome interval associated with resistance. Within these intervals, we identified candidate genes, including wall-associated kinases similar to *Stb6*, the first cloned STB resistance gene. Our analysis revealed a strong negative genetic correlation between tolerance and resistance to STB, indicative of a trade-off. Such a trade-off between tolerance and resistance would hinder breeding simultaneously for both traits, but our findings suggest a way forward using marker-assisted breeding. We expect that the methods described here can be used to characterize tolerance to other fungal diseases that produce visible fruiting bodies, such as speckled leaf blotch on barley, potentially unveiling conserved tolerance mechanisms shared among plant species.

## Introduction

Plants defend themselves against pathogens using either resistance, measured as the host’s ability to limit pathogen multiplication, or tolerance, measured as the host’s ability to reduce the negative effects of infection (Pagan and Garcia-Arenal 2020). A fundamental difference between these two strategies is that resistance reduces the multiplication rate of the pathogen, whereas tolerance does not. Tolerance was first recognized by plant pathologists in 1894 (Cobb 1894), and it is thought to be a host defense strategy as common and important as resistance (Pagan and Garcia-Arenal 2020). Yet, our knowledge of the mechanisms and genes controlling tolerance pales in comparison to our knowledge of the mechanisms and genes underlying resistance. This stems in part from the difficulty of measuring tolerance in plants and also from a lack of agreement in how tolerance should be defined. The excellent, comprehensive review by Pagan and Garcia-Arenal (2020) presents well-reasoned definitions that should resolve the latter difficulty.

The genetic basis of tolerance in plants has rarely been studied and remains poorly understood. The Pagan and Garcia-Arenal (2020) review identified ten examples where tolerance was inferred and its genetic basis was analyzed. In most of these cases, tolerance appeared to be a quantitative trait, involving from one to 70 quantitative trait loci (QTL) or candidate genes, but in some of these cases, it remains unclear whether the measured trait was truly tolerance or a form of resistance that was treated as tolerance (Ayala et al., 2002; Williams et al., 2003; Han et al., 2008). For example, Han et al. (2008) identified eight QTLs associated with tolerance to *Phytophthora sojae* in soybeans, using the proportion of surviving plants as a proxy for tolerance. But it is possible that the plants surviving an exposure to *P. sojae* were displaying partial resistance instead of tolerance to this pathogen. The most extensive work on tolerance has been conducted with plant viruses, including a study where tolerance to the barley yellow dwarf virus in wheat was ascribed to 22 QTLs of minor effect (Ayala et al. 2002). In tomatoes, a MAP kinase was found to enhance tolerance to the tomato yellow leaf curl virus (TYLCV) by regulating salicylic acid and jasmonic acid signaling (Li et al. 2017). More recently, the overexpression of a cellulose synthase-like gene was shown to boost tolerance to TYLCV (Choe et al., 2021). Functional alleles of flowering repressor genes in *Arabidopsis thaliana* were found to contribute to plant tolerance to cucumber mosaic virus (Shukla et al., 2021). Tamisier et al. (2022) identified candidate genes for potato virus Y tolerance in peppers (*Capsicum annuum*), including a cluster of NBS-LRR genes. But we are not aware of any studies that have identified candidate genes specifically associated with tolerance to fungal plant pathogens until now.

Septoria tritici blotch (STB) is the most damaging disease of wheat in Europe (Jorgensen et al. 2014) and among the most important diseases of wheat globally (Savary et al. 2019). STB is caused by the fungus *Zymoseptoria tritici*, a pathogen that has co-evolved with wheat for more than 10,000 years (Stukenbrock et al. 2007) and has a high evolutionary potential (McDonald et al. 2022). The most common strategies for controlling STB are deployments of fungicides and STB-resistant wheat cultivars. Exposed *Z. tritici* populations typically evolve resistance to fungicides and virulence against resistant cultivars within a few years of deployment (McDonald and Mundt 2016; McDonald et al. 2019; Kildea et al. 2020). The genetic basis of fungicide resistance and virulence has been explored in several populations of *Z. tritici*, leading to the discovery, cloning and functional validation of several of the underlying genes for both traits (Meile et al. 2018; Zhong et al. 2017; Amezrou et al. 2023; Garnault et al. 2019). Several STB resistance genes in wheat have also been cloned and functionally validated (Saintenac et al. 2021; Saintenac et al. 2018; Hafeez et al. 2023). The cloned pathogen avirulence genes and cloned wheat resistance genes have been shown to largely conform to the gene-for-gene concept of plant-pathogen interactions, though resistance to STB does not appear to involve a hypersensitive response (Saintenac et al. 2018).

Leaf-level tolerance to *Z. tritici* is a quantitative trait that was recently discovered in wheat (Mikaberidze and McDonald 2020) by using automated image analyses that could accurately quantify the leaf area affected by STB and count the number of fungal fruiting bodies (called pycnidia) found on the same leaf (Stewart et al. 2016; Karisto et al. 2018). These measures were then used to quantify degrees of STB resistance and STB tolerance in wheat, the latter using a novel measure called kappa, in 335 elite winter wheat cultivars growing in the same field. Resistance was quantified as the average number of pycnidia on a leaf, *N*_p_, adjusted for the overall leaf area. More resistant plant genotypes suppressed pathogen reproduction, and consequently carried fewer pycnidia on their leaves. Kappa is an exponential slope that characterizes the negative relationship between green leaf area and the number of pycnidia on a leaf. Lower kappa values correspond to higher tolerance: when inhabited by pathogen populations of the same size (i.e., when leaves carry the same numbers of pycnidia), more tolerant plant genotypes retain larger green leaf areas than less tolerant genotypes (Figure 1). Mikaberidze & McDonald (2020) showed that there was a wide, continuous variation in both resistance and leaf tolerance across the 335 wheat cultivars. They also found a negative relationship between tolerance and resistance, indicative of a trade-off between these traits.

**Figure 1.**
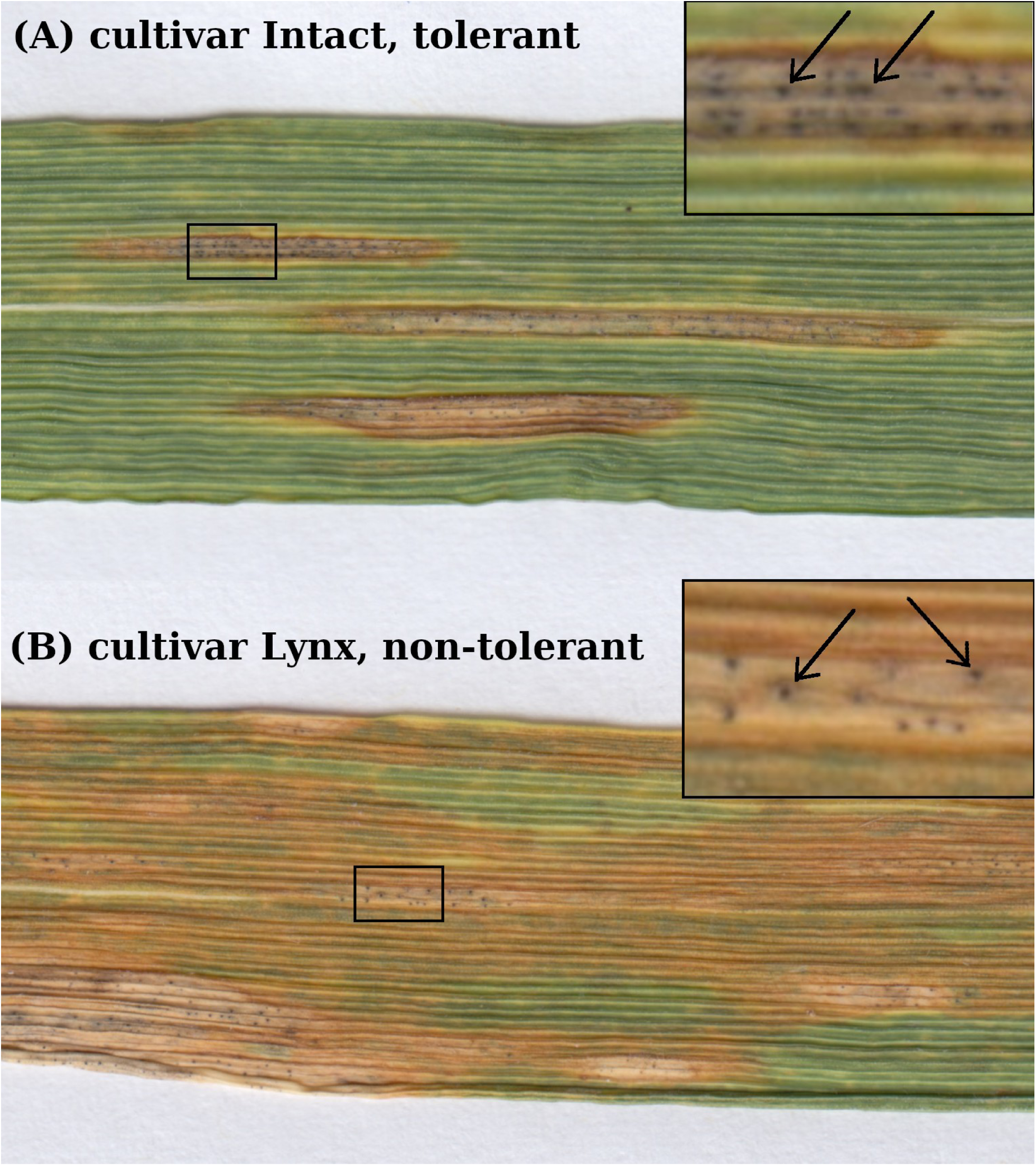
Illustration of leaf tolerance. STB symptoms on wheat leaves can be seen as characteristic necrotic lesions with pycnidia (seen as small, black, round structures within lesions) on cultivar Intact (A, tolerant) vs cultivar Lynx (B, non-tolerant). The number of pycnidia is similar in the two images, but the non-tolerant leaf has a larger disease-induced necrotic area (and a smaller green leaf area), and therefore suffers more damage from disease compared to the tolerant leaf. Black rectangles are 4x magnified in the insets, where two characteristic pycnidia are indicated by arrows. To quantify leaf tolerance it is not sufficient to compare two leaves, instead, we analyzed large numbers of diseased leaves from each cultivar and estimated tolerance as the negative exponential slope of how green leaf areas decrease versus the number of pycnidia on a leaf (see Figure 4 and Mikaberidze & McDonald 2020).

Before the discovery of leaf tolerance, we used these data to conduct a genome-wide association study (GWAS) to analyze the genetic architecture of STB resistance in the 335 wheat cultivars and identified several chromosome regions that contained interesting candidate STB resistance genes (Yates et al., 2019). Here we use the leaf tolerance trait kappa to conduct a GWAS aiming to elucidate the genetic architecture of tolerance and identify candidate genes that may be associated with leaf tolerance. We also sought to determine if the trade-off between tolerance and resistance to STB has a genetic basis. We discovered that the genetic associations for tolerance were independent from the previously described genetic associations for resistance to this pathogen. We identified four chromosome intervals associated with leaf-level tolerance and a separate chromosome interval associated with resistance. A bivariate GWAS that jointly considered both tolerance and resistance did not provide significant marker associations, even though there was a significant negative genetic correlation between these traits. Within each of the significant chromosome intervals, we identified candidate genes associated with tolerance.

## Results

Leaf tolerance (quantified as kappa) and resistance to STB (quantified as the number of pycnidia per leaf, *N*_p_) were measured based on the analysis of 11,152 individual images of naturally infected leaves coming from 335 elite winter wheat cultivars growing in a replicated field experiment (Karisto et al. 2018). On average, each leaf was infected by a different strain of *Z. tritici* (Lorrain et al. 2024; McDonald et al. 2022), hence the measures of leaf tolerance and resistance calculated for each cultivar represent average values across a very large number of pathogen strains. Despite the large variance associated with infections involving thousands of pathogen strains, the heritabilities were high for both leaf tolerance (0.44 for kappa) and resistance (0.88 for *N*_p_). There were strong phenotypic (*r_p_* = −0.40, *p*<0.001) and genetic (*r_g_* = −0.67, *p*<0.001; Figure 2) correlations between leaf tolerance and resistance.

**Figure 2.**
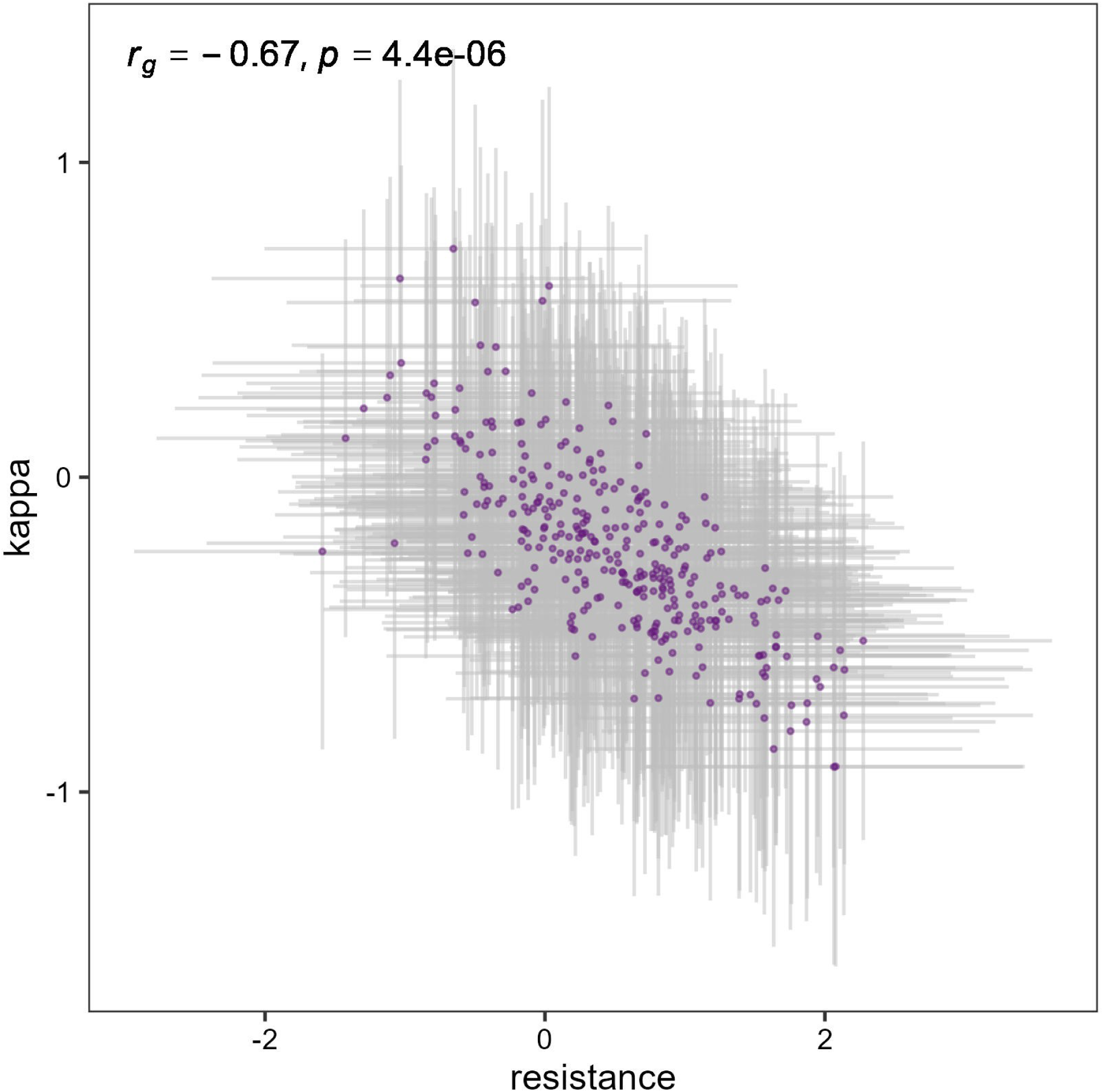
Genetic correlation between tolerance (quantified as kappa) and resistance (quantified as *N*_p_). Purple dots represent the empirical best linear unbiased predictors extracted from the bivariate model (Eq. 3) and gray lines represent the standard deviations.

The leaf tolerance and resistance phenotypes were used to conduct a GWAS that included, after filtering, 9,125 single nucleotide polymorphisms (SNPs) in 330 of the wheat cultivars. The GWAS identified three marker-trait associations (MTAs) for leaf tolerance that exceeded the Bonferroni significance threshold, located on chromosomes 2D, 6B and 7D (Figure 3). An additional leaf tolerance MTA on chromosome 7A fell just below the Bonferroni threshold (LOD = 5.19, Figure 3), but was included in further analyses. For resistance, *N*_p_, one significant MTA was detected on chromosome 5A.

**Figure 3.**
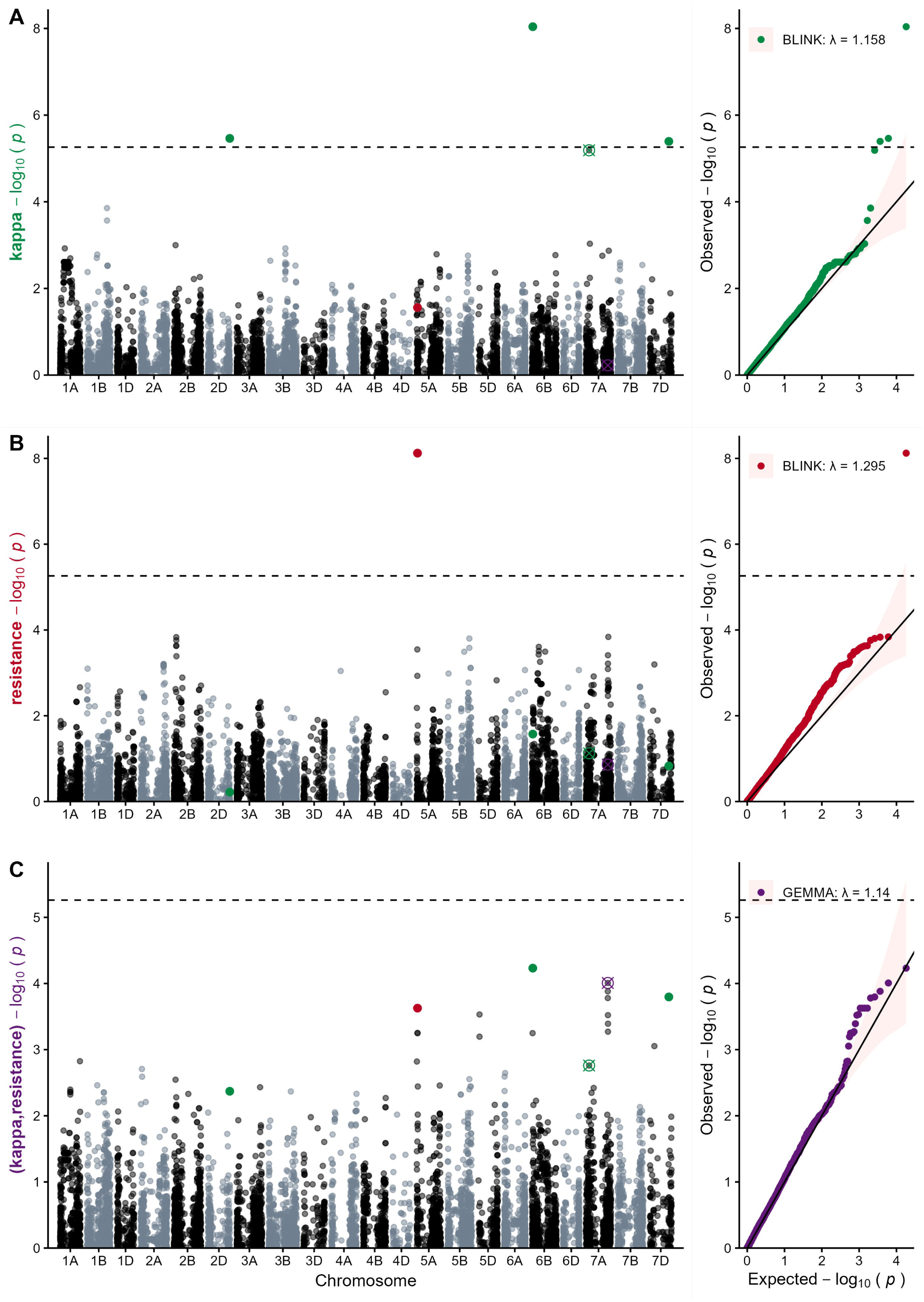
Manhattan and QQ-plots depicting the GWAS results for tolerance (kappa) (A), resistance (B) and the combined, bivariate GWAS (C). Green, red and purple dots in the Manhattan plots represent the SNPs significantly associated with the respective traits: kappa, resistance and the bivariate model combining kappa and resistance. Crossed circles represent marker-trait associations that were not significant at the Bonferonni threshold, but were nevertheless considered in additional analyses. The legends in the QQ-plots indicate the applied GWAS model and the genome-wide inflation coefficient λ.

The bivariate GWAS that jointly considered leaf tolerance and resistance did not detect any significant associations, though the MTAs observed on chromosomes 5A, 6B and 7D in the univariate analyses remained distinct relative to other SNPs (Figure 3, panel C). Though it was not significant at the Bonferroni threshold, a distinctive peak visible on chromosome 7A (LOD = 4.01, Figure 3, panel C) was included in further analyses.

The SNPs and chromosome positions for each of the six identified MTAs are shown in Table 1. For each MTA, a chromosome interval was defined by including the DNA sequences positioned 2.5 million base pairs (Mbp) in each direction from the significant SNP, i.e., by placing the most significant SNP at the center of a 5 Mbp interval on the IWGSC reference sequence v1.0 (IWGSC, 2018). The positions of these chromosome intervals were then compared to the positions of chromosome intervals containing STB resistance QTL that were reported in recent publications (Yates et al. 2019; Zakieh et al. 2023; Mekonnen et al. 2021; Alemu et al. 2021; Mahboubi et al. 2022) to determine if there were any overlaps (Supplementary Table S1). The leaf tolerance MTA on chromosome 2D was found to be 0.9 Mbp upstream of an STB resistance MTA that was previously detected in a greenhouse experiment where seedlings of 316 Nordic breeding lines were inoculated with two Nordic strains of *Z. tritici* (Zakieh et al. 2023). The resistance MTA on chromosome 5A was found to be within the same interval as a different STB resistance trait, PLACL (percentage of leaf area covered by lesions), that was identified in our earlier analyses of the same dataset (Yates et al. 2019). The tolerance MTA on chromosome 7D was found to be 1.6 Mbp downstream of the STB resistance QTL detected in a greenhouse experiment where seedlings of 185 wheat genotypes of globally diverse origin were inoculated with ten different *Z. tritici* strains of global origin (Mahboubi et al., 2022). Using the consensus map by Wang et al. (2014), we also compared our results to earlier publications reviewed in Brown et al. (2015). There, we found that the resistance MTA on chromosome 5A and the tolerance MTAs on chromosomes 6B, 7A and 7D were within the reported intervals for MQTL19, 21, 24 and 27, respectively. However, these intervals are too large (45, 22, 55 and 13 cM, respectively) to be confident that these overlaps are biologically meaningful. Therefore, while our tolerance MTA on chromosomes 2D, and 7D fall within previously reported resistance intervals, our tolerance MTA on chromosomes 6B and 7A are likely to constitute new loci or refined loci for tolerance within meta-QTL previously associated with resistance.

**Table 1:**
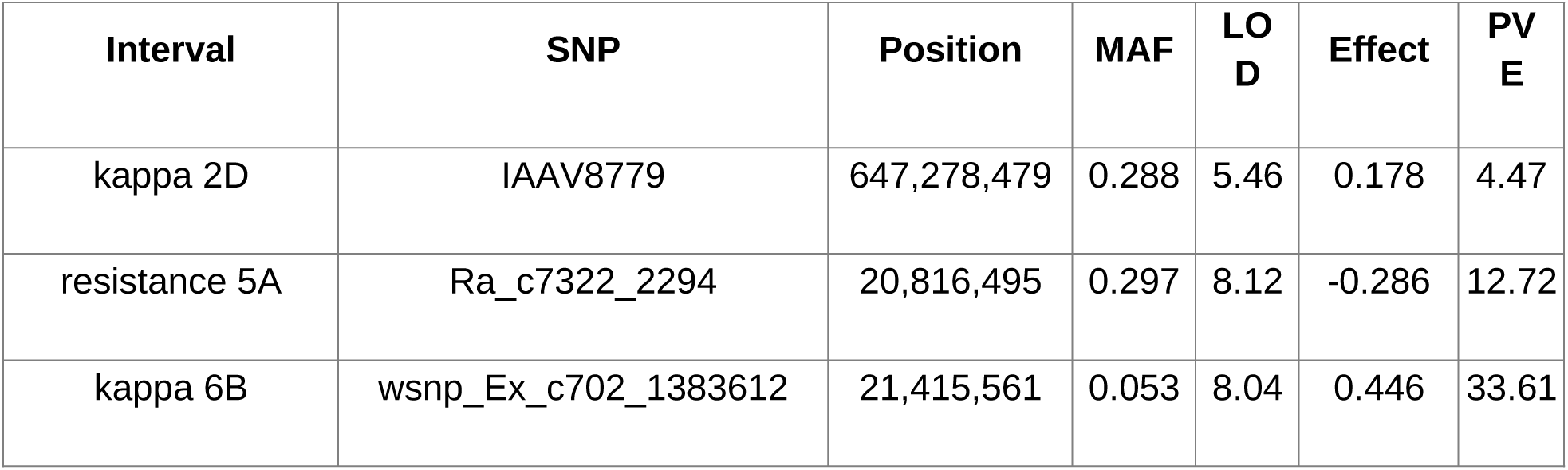
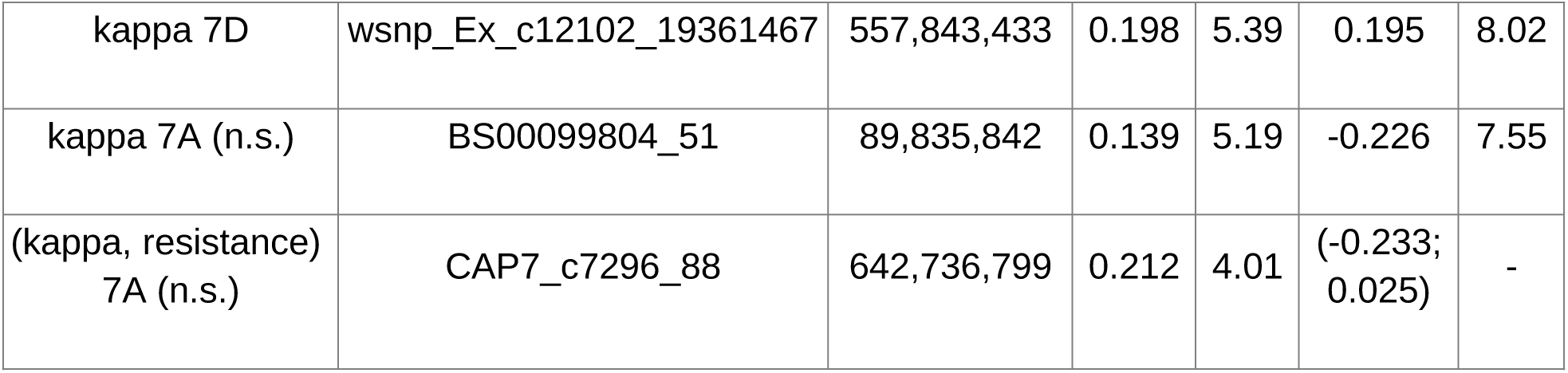
Marker-trait associations (MTAs) for tolerance (kappa), resistance and the combined, bivariate GWAS. For each MTA, we report the physical position, minor allele frequency (MAF), LOD score, the SNP effect estimates, and the percentage of phenotypic variance explained (PVE). Non-significant MTAs are indicated with “n.s.”

| Interval | SNP | Position | MAF | LOD | Effect | PVE |
| --- | --- | --- | --- | --- | --- | --- |
| kappa 2D | IAAV8779 | 647,278,479 | 0.288 | 5.46 | 0.178 | 4.47 |
| resistance 5A | Ra_c7322_2294 | 20,816,495 | 0.297 | 8.12 | -0.286 | 12.72 |
| kappa 6B | wsnp_Ex_c702_1383612 | 21,415,561 | 0.053 | 8.04 | 0.446 | 33.61 |
| kappa 7D | wsnp_Ex_c12102_19361467 | 557,843,433 | 0.198 | 5.39 | 0.195 | 8.02 |
| kappa 7A (n.s.) | BS00099804_51 | 89,835,842 | 0.139 | 5.19 | -0.226 | 7.55 |
| (kappa, resistance)<br>7A (n.s.) | CAP7_c7296_88 | 642,736,799 | 0.212 | 4.01 | (-0.233;<br>0.025) | - |

In order to identify candidate genes, we employed a two-pronged approach: (i) a gene motif overrepresentation analysis and (ii) a differential gene expression analysis based on published data (Ramírez-González et al., 2018, Rudd et al., 2015). To accomplish (i), we first searched the IWGSC refseq1.0 annotation (IWGSC, 2018) within 5 Mbp windows around each of the six identified MTAs. We selected candidate genes within these regions based on their functional description (Supplementary Tables S2-S7) and classified them into 13 motif groups. Next, we compared the occurrence of these groups within each interval to their occurrence in 10,000 intervals of 5 Mbp size randomly chosen across the genome (see Materials and Methods). This analysis allowed us to determine whether the 5 Mbp intervals around the MTAs we identified were more likely to contain a gene with a specific function related to plant defense as compared to intervals of the same size randomly chosen across the genome. Seven of these motif groups showed a significant (p<0.05) overrepresentation compared to random genome intervals and contained a total of 16 candidate genes (Table 2). For details regarding the motif groups and individual genes, see Supplementary Tables S8, S9 and S10.

**Table 2:** Overrepresentation of gene motifs within 5 Mbp intervals around marker-trait associations. The significance of motif overrepresentation was determined based on the probability distribution of their occurrence across 10,000 random samples. Column 4 shows the total occurrence of motifs across the entire genome. Columns 5-8 show the expected occurrence of motifs in 90%, 95%, 99% and 99.9% of the random samples, respectively. Column 9 shows the observed occurrence of motifs in the intervals around MTAs. The symbols ‘***’, ‘**’, ‘*’, and ‘.’ indicate the level of significance of overrepresentation with p<0.001, p<0.01, p<0.05 and p<0.1 respectively; ‘n.s.’ denotes no significant overrepresentation.

| interval | genes<br>in<br>interval | gene group | occurrence<br>genome | expected <= probability<br>quantile |  |  |  | observed |  |
| --- | --- | --- | --- | --- | --- | --- | --- | --- | --- |
|  |  |  |  | q90 | q95 | q99 | q99.9 |  |  |
| kappa<br>2D | 97 | Cysteine<br>proteinase<br>inhibitor | 101 | 0 | 0 | 3 | 6 | 4 | ** |
|  |  | GCN1 | 6 | 0 | 0 | 0 | 1 | 1 | ** |
|  |  | Cysteine<br>protease | 225 | 1 | 1 | 6 | 11 | 2 | * |
|  |  | LRR | 3215 | 16 | 20 | 36 | 50 | 17 | . |
|  |  | Expansin | 262 | 1 | 2 | 5 | 10 | 1 | n.s. |
|  |  | Wat1 | 194 | 1 | 2 | 3 | 6 | 1 | n.s. |
| kappa<br>6B | 62 | LRR | 3215 | 16 | 20 | 36 | 50 | 17 | . |
|  |  | Wall-associated<br>kinase | 95 | 0 | 1 | 2 | 4 | 1 | . |
| kappa<br>7A (n.s.) | 43 | DCD domain | 34 | 0 | 0 | 1 | 2 | 1 | * |
|  |  | Lectin RLK | 209 | 1 | 1 | 3 | 4 | 1 | n.s. |
| kappa<br>7D | 30 | Ran-binding | 12 | 0 | 0 | 0 | 0 | 1 | *** |
|  |  | Aspartate--<br>tRNA ligase | 16 | 0 | 0 | 1 | 1 | 1 | * |
|  |  | Wound-responsive | 69 | 0 | 0 | 3 | 14 | 2 | * |
| resistance<br>5A | 50 | Cysteine<br>protease | 225 | 1 | 1 | 6 | 11 | 1 | n.s. |
|  |  | LRR | 3215 | 16 | 20 | 36 | 50 | 8 | n.s. |
| (kappa,<br>resistance)<br>7A (n.s.) | 61 | Ran-binding | 12 | 0 | 0 | 0 | 0 | 1 | *** |
|  |  | Wound-responsive | 69 | 0 | 0 | 3 | 14 | 3 | * |
|  |  | LRR | 3215 | 16 | 20 | 36 | 50 | 1 | n.s. |

Around the MTAs for leaf tolerance, we found a significant overrepresentation of gene motifs that can be associated with programmed cell death, including, cysteine proteinase inhibitors (observed=4, p<0.01), cysteine proteases (observed=2, p<0.05), DCD domain (observed=1, p<0.05) and wound responsive motifs (observed=2, p<0.05). We also found a significant overrepresentation of disease resistance motifs (GCN1, observed=1, p<0.01), aspartate--tRNA ligase (observed=1, p<0.05), cell wall-associated Ran-binding (observed=1, p<0.001), and wall-associated kinases (WAK; observed=1, with a more relaxed significance threshold p<0.1). The bivariate MTA on chromosome 7A also showed a significant overrepresentation of cell wall associated Ran-binding (observed=1, p<0.001) and wound-responsive (observed=2, p<0.05) motifs. Around the resistance MTA on chromosome 5A, we observed one cysteine protease and eight genes with “leucine rich repeat” (LRR) motifs, neither of which was significantly overrepresented.

We note that based on the Automated Assignment of Human Readable Descriptions (AHRD), only one WAK gene has been found in the tolerance MTA interval on chromosome 6B, but we examined this interval more thoroughly (based on InterPro; https://www.ebi.ac.uk/interpro/) and found three more WAKs (which have been identified as receptor-like kinases by AHRD). We consider these four WAKs in this interval to be interesting because *STB6,* the first cloned resistance gene for STB, is also a wall-associated serine/threonine kinase with a galacturonan-binding domain (Saintenac et al. 2018); although the four WAKs have an epidermal growth factor (EGF) domain, which *STB6* does not have.

To accomplish (ii), we compared gene expression across all our MTA intervals between *Z. tritici*-infected seedlings and mock-inoculated controls using published data (Rudd et al., 2015; Ramírez-González et al, 2018; data available via www.wheat-expression.com) and identified candidate genes as those having significant differential expression. We found that 26 genes around the tolerance MTAs, seven genes around the resistance MTA and six genes around the MTA for the bivariate model were significantly differentially expressed (Table 3; Supplementary Table S11 for more details). Among the MTAs for leaf tolerance, we found significant differences in gene expression for three glutathione S-transferases (log2-fold difference ranging from 3.7 to 8.3), a receptor-like protein kinase (log2-fold difference of 1.6), a transposon protein with a NAC transcription domain (log2-fold difference of 5.5), and a RING finger protein that was down-regulated (log2-fold difference of −1.6). Another glutathione S transferase was significantly up-regulated (log2-fold difference of 3.8) in the MTA for resistance.

**Table 3:**
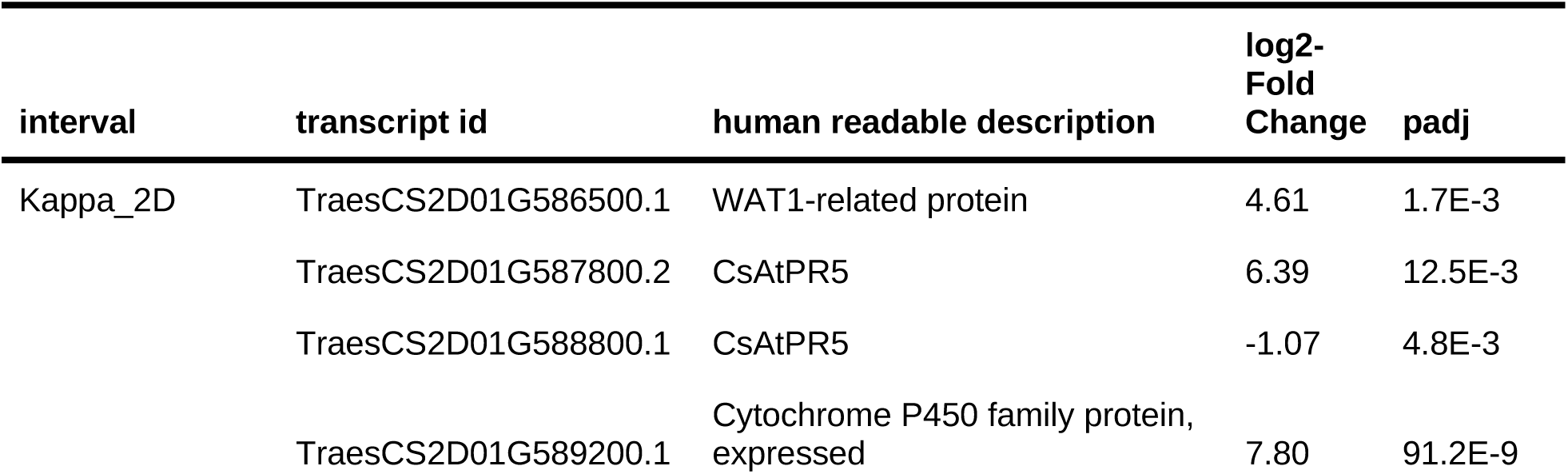
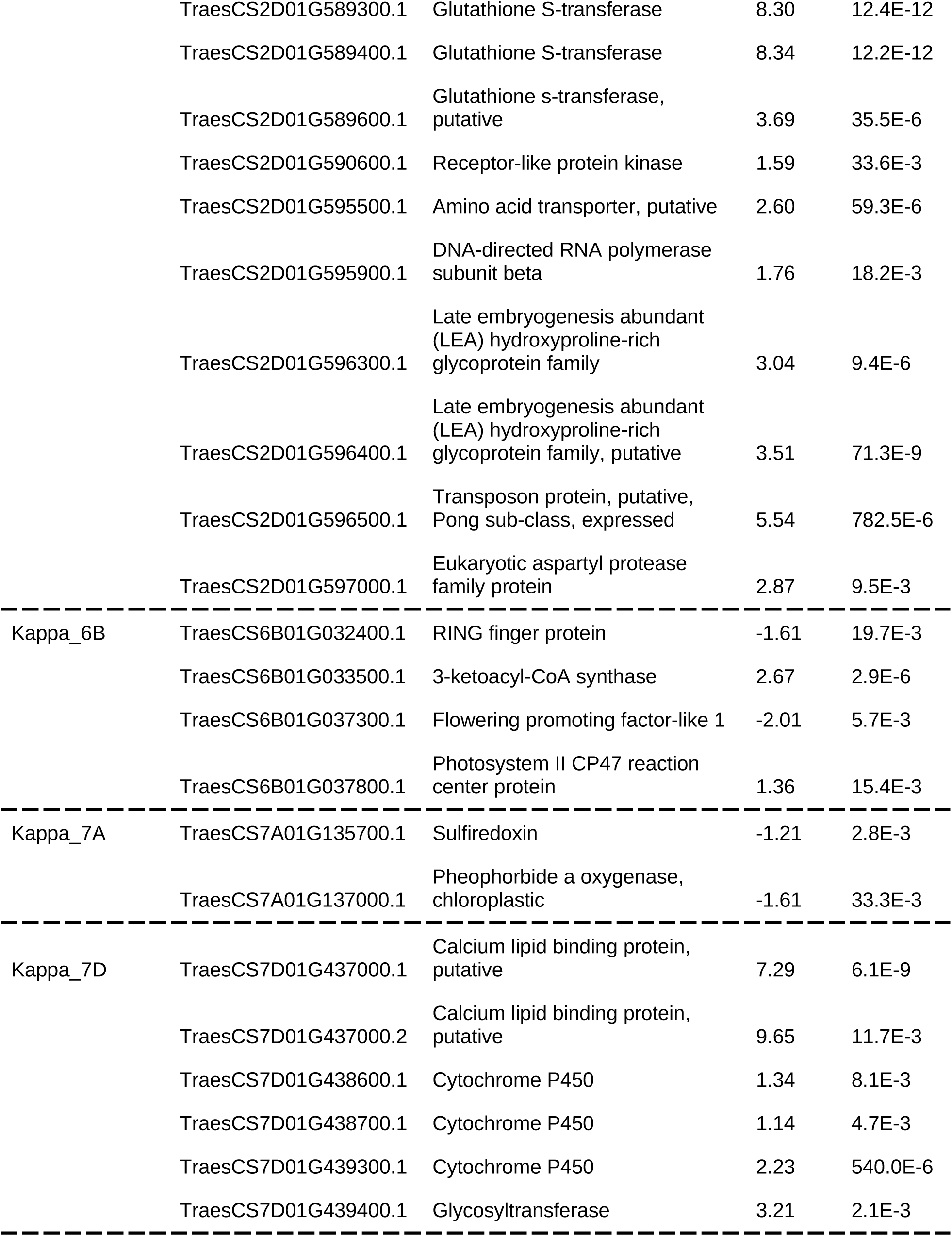
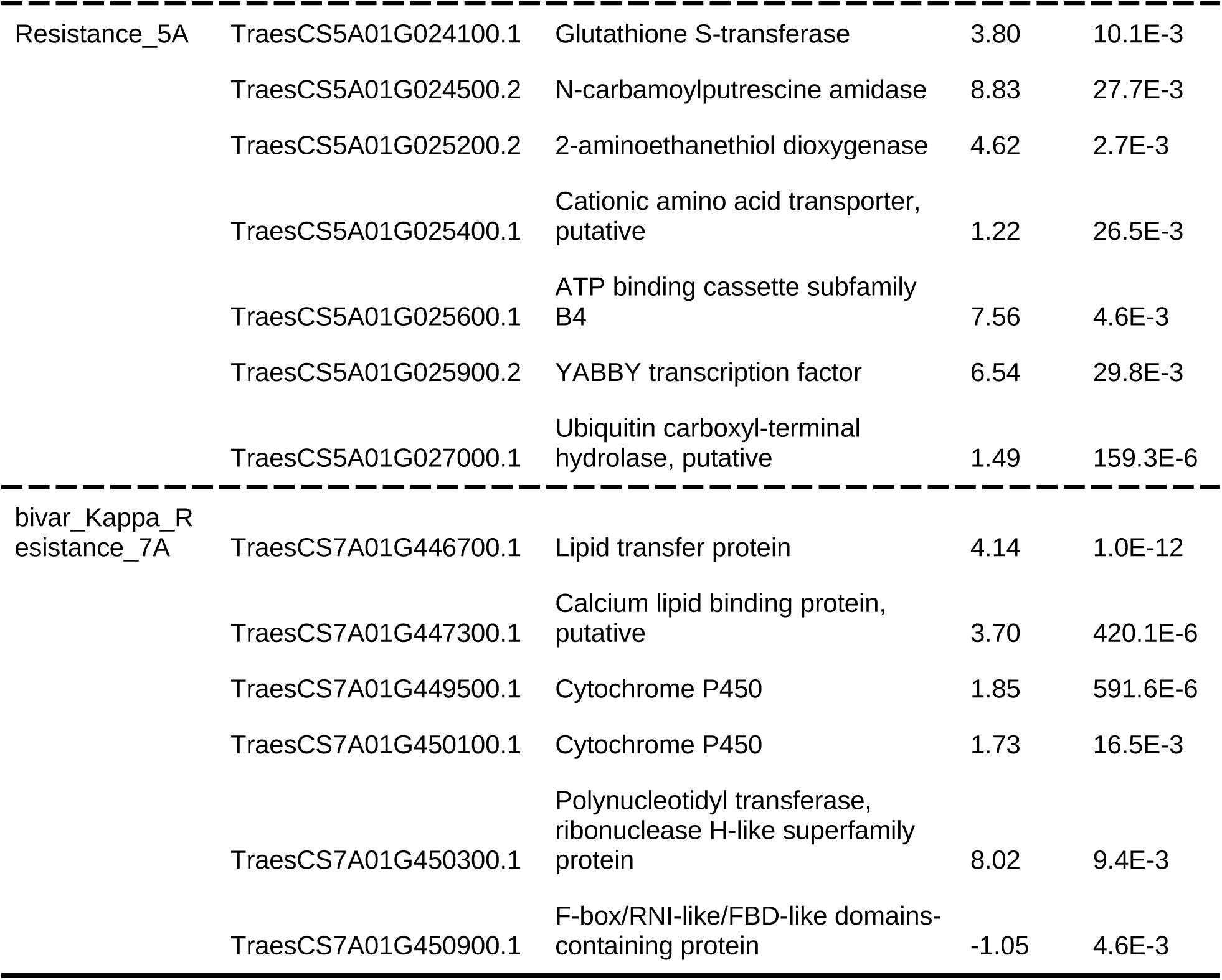
Differentially expressed genes within 5 Mbp intervals around marker-trait associations. Transcript id, start and end denote the isoform and position (bp), distance denotes the distance (bp) to the respective marker-trait association, baseMean denotes the average of the normalized count values, lfcSE is the standard error of the log2-fold change in expression (log2FoldChange) and padj is the adjusted p-value.

## Discussion

Though several studies have analyzed the genetics of disease tolerance in plants (Pagan & Garcia Arenal 2020), we believe this is the first study to reveal the genetic basis of tolerance to a fungal plant pathogen and to identify candidate genes that may confer the leaf tolerance phenotype. Among these candidates are wall-associated kinases with galacturonan-binding and serine/threonine kinase domains similar to *Stb6*, the first cloned STB resistance gene (Saintenac et al. 2018). Our automated analyses of more than 11,000 scanned leaves revealed that wheat’s response to an STB infection can be dissected into resistance and tolerance components that display high and moderate heritabilities, respectively. In the analyzed population of 330 elite winter wheat cultivars, the two traits showed a strongly negative genetic correlation (Figure 2), supporting our previous report of a substantial negative phenotypic correlation indicative of a trade-off between tolerance and resistance (Mikaberidze & McDonald 2020).

While screening for germplasm that performs well during an STB epidemic, wheat breeders cannot distinguish between the leaf tolerance and resistance traits using traditional visual scoring. A more detailed phenotyping based on leaf image analysis is required to distinguish between them. As a result, breeding based on visual assessments may select germplasm that has higher resistance or higher tolerance, both of which could produce an improved yield response to STB infections compared to STB susceptible germplasm. This may explain why the STB tolerance QTL that we identified on chromosome 2A overlapped with an STB resistance QTL identified in Nordic breeding lines based on traditional visual scoring of STB under greenhouse conditions (Supplementary Table S1; Zakieh et al., 2023). Furthermore, because breeders have been selecting for better performance in elite European winter wheat over several decades using visual scoring, we would expect these efforts to result in cultivars with high values for both tolerance and resistance. But it was rare to find elite winter wheat cultivars that show high values for both traits (Figure 2), suggesting that there is indeed a trade-off between the two traits. These interpretations are consistent with our previous analysis, where we found a positive correlation between cultivar release year and degree of tolerance and an absence of such a relationship for resistance, which suggested that European wheat breeders may have been selecting for tolerance instead of resistance to STB during recent decades (Mikaberidze & McDonald 2020).

Should wheat breeding programs seek to limit STB damage by combining STB resistance and tolerance into the same cultivar? This would require that the two traits are encoded by a shared pathway or by separate sets of genes that can be recombined into the same lineage, as well as the absence of epistasis. Our analyses reveal a nuanced picture of genetic connections between STB resistance and tolerance. The significant and strongly negative genetic correlation between the two traits suggests that the traits are not independent of each other. Yet the univariate GWAS analyses identified different chromosome intervals associated with tolerance and resistance, while the bivariate GWAS revealed only one association that was visually striking, but not statistically significant. A similar pattern was found in studies of resistance and tolerance of pepper plants to potato virus Y: the two traits exhibited significant negative phenotypic and genetic correlations, the GWAS identified markers that were significantly associated with either tolerance or resistance, but none of the markers were shared between the two traits (Tamisier et al., 2020; Tamisier et al., 2022). A possible explanation of these outcomes is that there could be several molecular pathways contributing to tolerance. Some of the pathways exhibit a negative genetic correlation with resistance and are underpinned by a large number of genes of small effect. Here, a negative correlation between tolerance and resistance may result from these genes exhibiting negative pleiotropy, whereby the same gene contributes to an increase in tolerance, but a decrease in resistance (or vice versa). Alternatively, a negative correlation between the two traits can be caused by linked monotropic genes: among several linked genes, some contribute to an increase in tolerance, while others contribute to a decrease in resistance (Gardner and Latta, 2007). Under either of these scenarios, we would not be able to capture these genes via MTAs in the GWAS. Other tolerance pathways could be independent of resistance, and conferred by fewer genes with larger effects. These would be identified as significant MTAs for tolerance in the GWAS, none of which were significantly associated with resistance. Hence, purely phenotypic selection for tolerance may inadvertently select against resistance and vice versa. However, marker-assisted selection for components of tolerance that are independent of resistance could avoid this pitfall and select for tolerance without compromising resistance. Additional experiments will be needed to further validate the tolerance MTAs and the associated candidate genes to enable this approach.

We identified tolerance candidate genes using analyses of both motif enrichment and differential gene expression. The enrichment analysis revealed a significant overrepresentation of genes encoding programmed cell death, leucine-rich repeats, responses to wounding, and wall-associated kinases. Many of these candidate genes encode functions that are typically associated with disease resistance, such as NBS-LRRs, lectin receptor-like kinases, and wall associated kinases, including genes that are similar to *Stb6*, the first cloned STB resistance gene (Saintenac et al. 2018). Similarly, Tamisier et al (2022) found a cluster of candidate NBS-LRR genes to be associated with tolerance of pepper plants to potato virus Y. The differential gene expression analysis also revealed genes known to be associated with disease resistance, including a receptor-like protein kinase. These findings suggest that there may be a common genetic architecture underlying plant resistance and tolerance. In particular the same pathogen sensing, recognition, and signaling processes may be involved at the beginning of the tolerance and resistance response pathways. Under this scenario, the same gene may confer tolerance to one pathogen, but resistance to a different pathogen. For example, a candidate NBS-LRR gene for potato virus Y tolerance in pepper (*Capsicum annuum*) shares 87.1% nucleotide identity with the *Bs2* gene in a different pepper species (*Capsicum chacoense*) known to confer resistance to bacterial spot disease (Tamisier et al., 2022). For tolerance, we found a greater overrepresentation of gene motifs associated with wound responses and programmed cell death (PCD) compared to the resistance trait (Table 2). We speculate that inhibition or appropriate regulation of wound response or PCD pathways may lead to increased leaf tolerance via reduction of excessive necrosis of leaf tissue.

While this is the first work to identify candidate genes affecting tolerance to fungal pathogens, we expect that the methods described in this paper can be applied to many other fungal diseases that produce visible fruiting bodies, such as speckled leaf blotch on barley, septoria leaf blotch on oats, or septoria leaf spot of tomatoes. As tolerance is analyzed in other plant pathosystems, it will become possible to compare candidate genes identified across different systems to determine if tolerance, like resistance, is encoded by conserved mechanisms shared among many plant species.

## Materials and Methods

Naturally infected penultimate leaves from 335 elite winter wheat cultivars were sampled from replicated plots during the 2016 field season as described in earlier publications (Karisto et al., 2018; Yates et al., 2019; Mikaberidze and McDonald, 2020). We estimate that at least half a million *Z. tritici* genotypes were present in the sampled plots (McDonald et al. 2022; Lorrain et al. 2024), thus our results are relevant for epidemics caused by highly diverse, natural pathogen populations. On average, 16 infected leaves from each plot were imaged using a flatbed scanner (Canon CanoScan LiDe 220) at 1200 dpi resolution to obtain the percentage of leaf area covered by lesions (PLACL, a damage function that reduces the green leaf area, a measure of disease-induced reduction of plant fitness for each leaf) and numbers of pycnidia (*N*_p_, associated with pathogen reproduction, a measure of the pathogen burden in each leaf) (Stewart et al., 2016; Karisto et al., 2018). In order to control for the effect of total leaf area on the number of pycnidia per leaf, we performed the adjustment *N*_p,*i*_ → (*A*_tot_/*A*_tot,*i*_) *N*_p,*i*_, where *N*_p,*i*_ and *A*_tot,*i*_ are the number of pycnidia and the total area of an individual leaf *i*, and *A*_tot_ is the mean total leaf area averaged over the entire dataset (Mikaberidze and McDonald, 2020).

These measures were then used to quantify degrees of STB resistance and tolerance, the latter using a novel measure called kappa. Kappa is an exponential slope that characterizes the negative relationship between green leaf area and the number of pycnidia on each leaf (Figure 4; Mikaberidze & McDonald, 2020). Small values of kappa indicate a high level of leaf tolerance, while large values of kappa indicate a low level of leaf tolerance. The kappa values were calculated for each cultivar based on measurements from approximately 32 leaves (each cultivar was replicated twice in the experiment), with each leaf typically infected by a different pathogen strain (McDonald et al. 2022; Lorrain et al. 2024). Hence the tolerance measures represented average values across a wide range of pathogen genotypes (≈32 *Z. tritici* strains) for each wheat cultivar and the range of kappa values encompassed a wide range of host genotypes (∼330 wheat cultivars). Resistance was quantified as the average number of pycnidia found on each leaf (*N*_p_; adjusted for the total leaf area). More susceptible cultivars allow higher numbers of pycnidia per leaf that translate to more pathogen reproduction, while more resistant cultivars limit the numbers of pycnidia per leaf and therefore reduce pathogen reproduction. Additional details on how tolerance and resistance were calculated for each cultivar can be found in Mikaberidze & McDonald 2020. The workflow for data acquisition and analysis is illustrated in Figure 4. The raw data stemming from the image analysis of each individual diseased leaf reported by Karisto et al. (2018) are available via Dryad Digital Repository: https://doi.org/10.5061/dryad.171q4. The processed phenotypic and genomic datasets underlying the outcomes of this study, and the code that can be used the reproduce the analyses are available via Zenodo https://doi.org/10.5281/zenodo.14962847.

**Figure 4.**
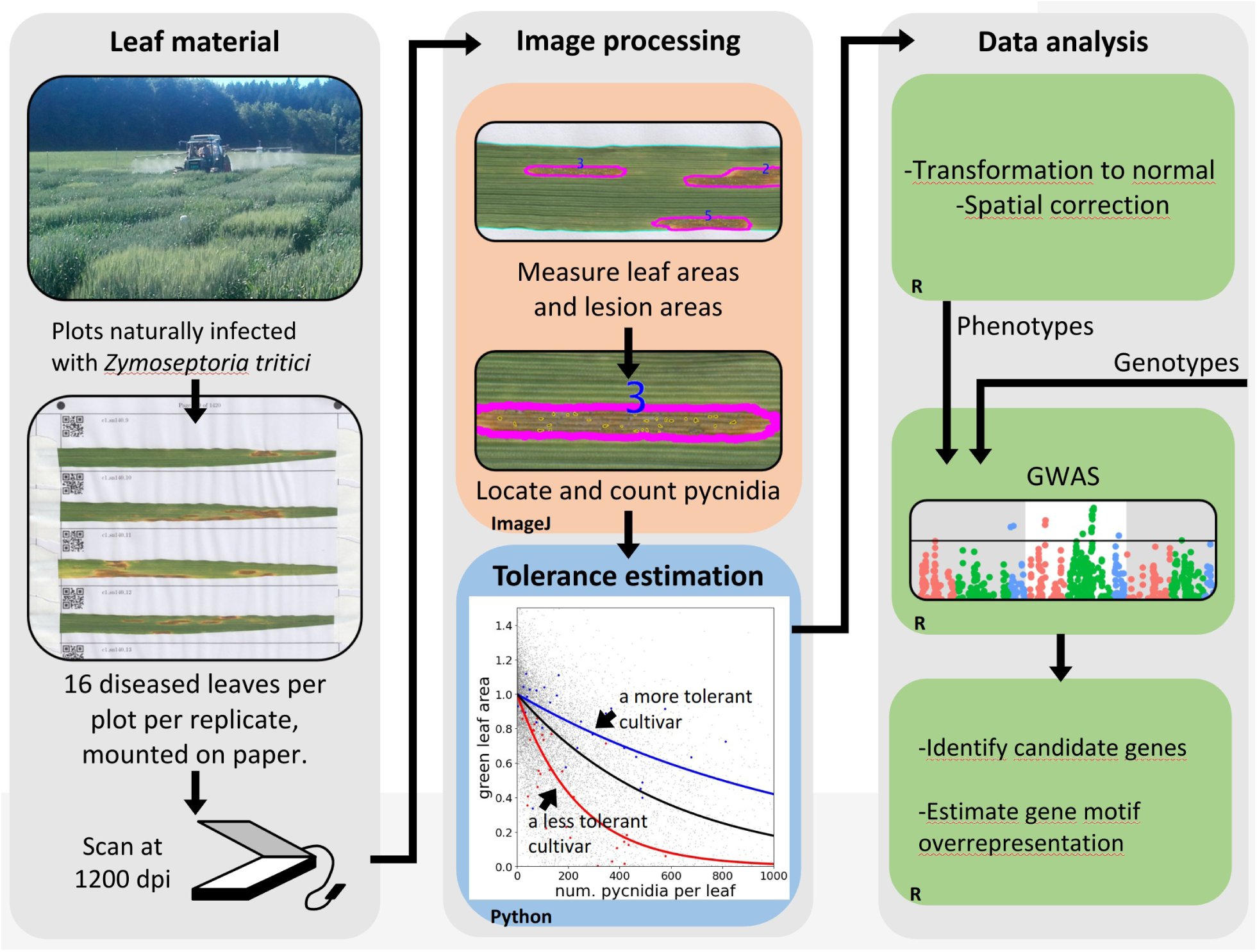
Workflow for the acquisition of phenotypic data, estimation of leaf tolerance and GWAS [modified from Figure 1 in (Yates et al., 2019) and Figure 3 in (Mikaberidze & McDonald, 2020].

### Statistical Analysis

An initial data inspection revealed strong skews in the distributions of both tolerance and resistance, which resulted in violations of the assumption of independence and normality of the residual distribution in the subsequent linear modeling (Supplementary Figure S1). For this reason, the raw data from individual plots was subjected to a rank-based inverse normal transformation (RINT) using the R-package RNOmni (v1.0.1.2; McCaw, 2023) before conducting further analyses. This brought the distributions of tolerance and resistance close to the normal distribution and resulted in independent, normally distributed residuals with a mean of zero.

To obtain best linear unbiased estimates (BLUEs) across replications while accounting for spatial variability, a spatial model using two-dimensional p-splines was fitted in the R-package SpATS (v1.0.18; Rodríguez-Álvarez *et al*., 2017). The two complete blocks were allocated diagonally in a virtual grid (see Kronenberg et al. 2021, Pérez-Valencia et al. 2022 for details), with rows and columns corresponding to the relative plot positions within each replicate of the experiment. The fitted model was:

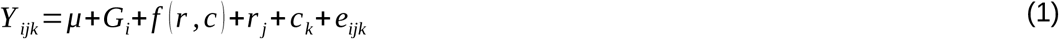

where *Y_ijk_* is the plot value of the respective trait: tolerance quantified as kappa or resistance quantified as *N*_p_. *μ* is the global intercept and *G_i_* is the response of the genotype *i*. To account for spatial variability, *r_j_* and *c_k_* represent the effects of the row *j* and column *k*, respectively, while *f(r,c)* is a smoothed bivariate surface across rows and columns within the virtual grid, thus fitting an independent spatial trend to each of the two replicates. From this model, the BLUEs were extracted to be used in the GWAS whereas spatially corrected plot values, comprising the BLUEs and residual errors but omitting spatial trends and other unwanted design factors, were used to calculate heritability and genetic correlations.

To calculate heritability, the model:

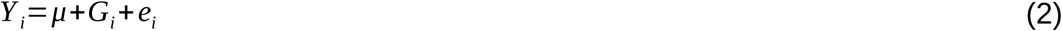

was fitted using the R-package asreml-R (v4.2.0.302, VSNi Team 2023), where *Y _i_* is the spatially corrected plot value from Eq. (1) for the respective trait, *G_i_*is the random genotype response with known variance-covariance structure based on the genome-wide identity by state (IBS) relationship matrix calculated from single nucleotide polymorphism (SNP) data using the R-package SNPRelate (v1.30.1; Zheng *et al*., 2012) and *e_i_* is the residual error. Heritability was estimated on a genotype difference basis: 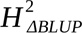 according to Eq. (24) in (Schmidt et al., 2019).

Phenotypic correlations were calculated as Pearson’s *r* based on the adjusted genotype means extracted from Eq. (1). To calculate the genetic correlation between kappa and resistance, Eq. (2) was expanded to the bivariate model:

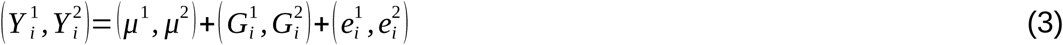

in the R-package asreml-R, where the superscripts 1 and 2 denote the two traits: kappa and *N*_p_, respectively. 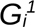 *and* 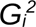 were again set as random with known variance-covariance structure based on IBS. The genetic correlation was then calculated from the estimated variance and covariance components from Eq. (3) following Holland et al. (2001):

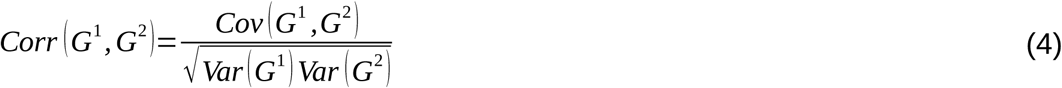

### Genome-wide association studies (GWAS)

Genome-wide association studies for the traits kappa, *N*_p_ and the combined, bivariate response were conducted following the same workflow as described by Roth & Kronenberg et al. (2024). Marker data were supplied by the GABI wheat consortium (Gogna *et al*., 2022) for the GABI genotypes and by the Agroscope wheat breeding program for the Swiss genotypes (Fossati and Brabant, 2003). Markers were mapped to the *Triticum aestivum* reference sequence v.1.0 (IWGSC et al., 2018) using ncbi-blast+ (v2.9.0-2). Equivocally mapped markers were excluded, and the remaining markers were filtered for a missing rate <0.05 and a minor allele frequency >0.05, resulting in 9,125 SNPs for 330 wheat cultivars. The remaining missing markers were imputed using *fastPHASE* (v.1.4; Scheet and Stephens, 2006) implemented in https://github.com/mwylerCH/HapMap_Imputation. Given the large size of the wheat genome (>14 Gbp), the SNP marker density is quite low, on average one marker per 1.6 Mbp, which may limit our capacity to detect marker-trait associations.

The univariate GWAS analyses for kappa and *N*_p_ were conducted using the Blink model (Huang *et al*., 2019) implemented in the R-package GAPIT3 (v.3.1.0) (Wang and Zhang, 2021). The bivariate GWAS was conducted using the software GEMMA (v0.98.1, Zhou and Stephens, 2014) using the first three principal components among SNP genotypes and the genome-wide IBS matrix to correct for population structure and relatedness, respectively.

Taking into account advances in GWAS methodologies in wheat since the publication of (Yates et al., 2019), we adapted our analyses in four ways. We used (i) an adjustment for spatial variability using the R-package SpATS (Rodríguez-Álvarez et al., 2017); (ii) a different transformation [RINT-transformation (McCaw et al., 2020) instead of log-transformation] that better satisfies the assumption of normality in the residuals of the applied linear models; (iii) a more comprehensive marker panel that includes Swiss cultivars, thus adding 11 genotypes previously excluded due to missing marker data; improved imputation of missing genotype data based on similarity of haplotype clusters around the missing genotype; and (iv) a different GWA model [single marker-based BLINK (Huang et al., 2019) instead of haplotype based PLINK (Purcell et al., 2007)] that better manages the systematic inflation of p-values in genetic association tests. To ensure reproducibility and better understand the effects of these four modifications, we reanalyzed the data of (Yates et al., 2019), and compared those results with the outcomes of our new GWAS pipeline applied to the same phenotypes. We illustrate the comparison of the two analyses in Supplemental Figure S2.

### Identification of candidate genes

To identify candidate genes, we searched the IWGSC refseq 1.0 annotation (IWGSC, 2018; https://urgi.versailles.inra.fr/download/iwgsc/) for 2.5 Mbp in each direction from the genome position of the associated SNP markers identified by GWAS. In a first step, likely candidates for either leaf tolerance or resistance were identified based on their functional description. Genes were categorized into motif groups based on their description (see Table 2; Supplementary Tables S8 & S9). Then the likelihood of occurrence for each motif was quantified using a bootstrapping approach. We examined 10,000 random 5 Mb intervals with a gene content >55 genes, i.e. the average of the gene content of the associated 5 Mb intervals, across the entire genome. We chose the size of the intervals to be 5 Mb, because this is below the characteristic LD decay distance (r2<0.2) for all the chromosomes of interest. In these intervals, we counted the occurrence of the selected candidate motifs identified in the intervals around the MTAs and calculated quantile distributions for the occurrence of the respective motifs across the random samples. A motif was considered significantly overrepresented if the occurrence in the identified interval was larger than the occurrence in 95% of the random intervals.

The leucine rich repeat (LRR) protein domain represents a characteristic feature of several classes of plant disease resistance proteins (Gururani et al., 2012). Hence, for the purpose of the representation analysis, we merged the gene groups “NBS-LRR”, “disease resistance proteins”, “RPM1“ and “RPP13” (all of which contain LRR) into a single “LRR” category (Table 2). We note that our overrepresentation analysis is based on Automated Assignment of Human Readable Descriptions (AHRD; https://github.com/groupschoof/AHRD?tab=readme-ov-file), which represents the official description of wheat genes published by the scientific community. However, there can be cases of misclassification, which lead to uncertainties in the occurrence of gene groups that are difficult to estimate, and for this reason the outcomes of this analysis need to be interpreted with caution.

Furthermore, differential gene expression analysis has been performed on the genes within the associated 5 Mbp intervals using publicly available transcript count data (Ramírez-González et al, 2018, https://www.wheat-expression.com/). The RNA-seq data originated from a gene expression study of *Z. tritici-*infected seedlings versus mock-infected seedlings (Rudd et al, 2015). There, seedlings of the wheat cultivar ‘Riband’ were infected with the *Z. tritici* isolate IPO323 and samples were taken at 5 different timepoints (1-21 days) after infection (see Rudd et al., 2015 for details). The outcomes of this analysis need to be interpreted with caution, because wheat gene expression can differ between controlled environments and the field environment, and also differ between seedlings and adult plants; gene expression can also be specific to host and pathogen genotypes. In this analysis, we pooled the transcript count data across timepoints. Differential gene expression was calculated across the whole genome using the R-package DEseq2 (v.1.44.0; Love et al., 2014). Genes were considered differentially expressed (DE) if their expression changed twofold and was significant (adjusted p-value <0.05). Only the DE genes within the 5 Mb intervals around each QTL were considered for candidate gene identification.

## Acknowledgements

We thank the group of Crop Science at ETH Zurich for allowing us to sample STB in their field experiment and in particular Andreas Hund for designing the field experiment and Hansueli Zellweger for crop husbandry. We are grateful to Petteri Karisto for leading the collection and processing of leaf samples, for help with preparing Figure 4, and for fruitful discussions, and to Steven Yates for conducting initial analyses. This work was supported by the Swiss National Science Foundation (SNSF) in the framework of the project DisPhenHiT (grant no. 206826). AM gratefully acknowledges financial support from the SNSF through the Ambizione grant PZ00P3_161453.

## Supplementary Material

Supplementary Figure S1: Residual vs. fitted values of the SpATS model, before and after rank based inverse normal transformation.

Supplementary Figure S2: Reproduction of Yates et al. (2019) GWAS outcomes using different models and marker sets.

Supplementary Table S1: Comparison of detected marker trait associations to previously reported QTL for *Z. tritici* resistance.

Supplementary Table S2: Gene list for the associated interval for kappa on chromosome 2D.

Supplementary Table S3: Gene list for the associated interval for kappa on chromosome 6B.

Supplementary Table S4: Gene list for the associated interval for kappa on chromosome 7A.

Supplementary Table S5: Gene list for the associated interval for kappa on chromosome 7D.

Supplementary Table S6: Gene list for the associated interval for resistance on chromosome 5A.

Supplementary Table S7: Gene list for the associated interval for the bivariate GWAS on chromosome 7A.

Supplementary Table S8: Gene motif group details including functional descriptions comprised in the motif groups, total occurrence of motifs across the genome, and gene IDs comprised in the motif groups.

Supplementary Table S9: Identified candidate gene motifs within 5 Mbp intervals around the respective marker-trait associations including candidate gene IDs per motif group.

Supplementary Table S10: Details on identified overrepresented genes including motif group, gene ID, start, end, functional description, Pfam description, interpro description, GO description and GO IDs from the IWGSC ref. seq. v1.0.

Supplementary Table S11: Details on differentially expressed genes within 5 Mbp intervals around marker-trait associations.

