## Supplementary figures S1 -- S2 for "A genome-wide association study identifies markers and candidate genes affecting tolerance to the wheat pathogen *Zymoseptoria tritici*"

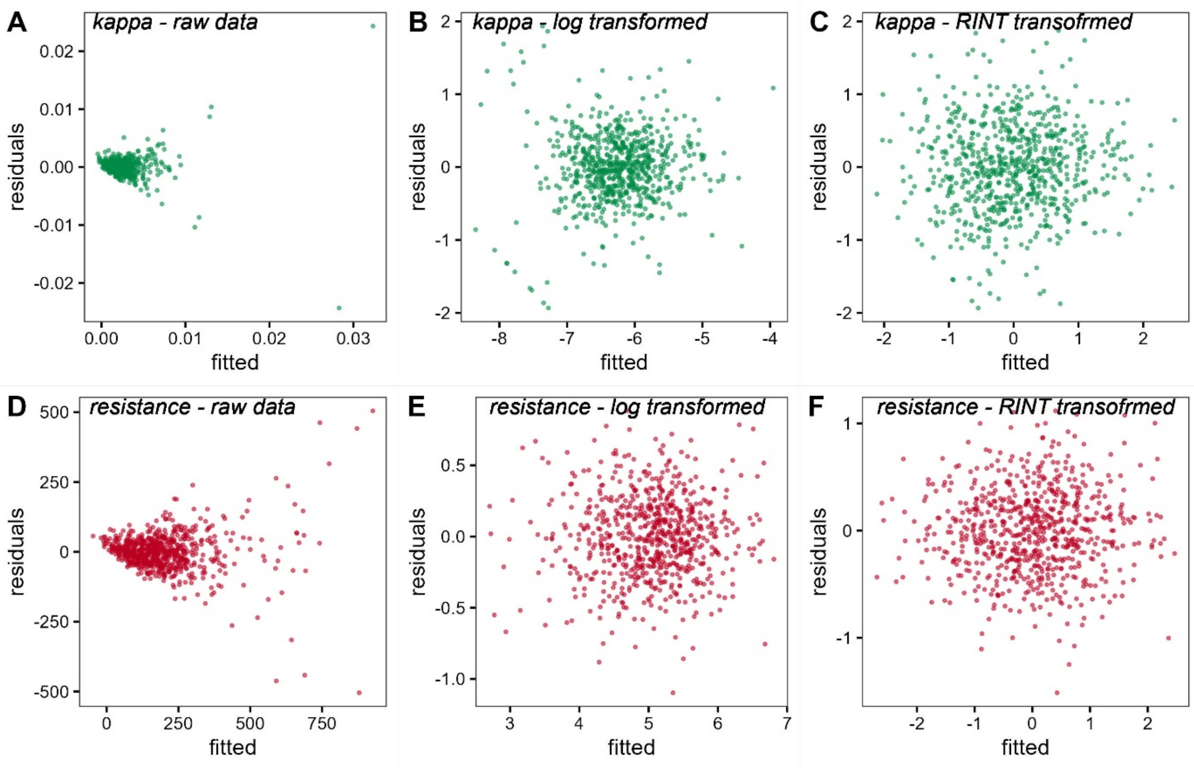


**Supplementary Figure S1: Scatter plots of residuals versus fitted values from the linear model applied to estimate best linear unbiased estimates from the phenotypic data (Equation 1).** A-C: Raw data, log-transformed data and rank-based inverse normal transformed (RINT) data for kappa (green dots); D-F: Raw data, log-transformed data and rank-based inverse normal transformed (RINT) data for resistance (red dots).

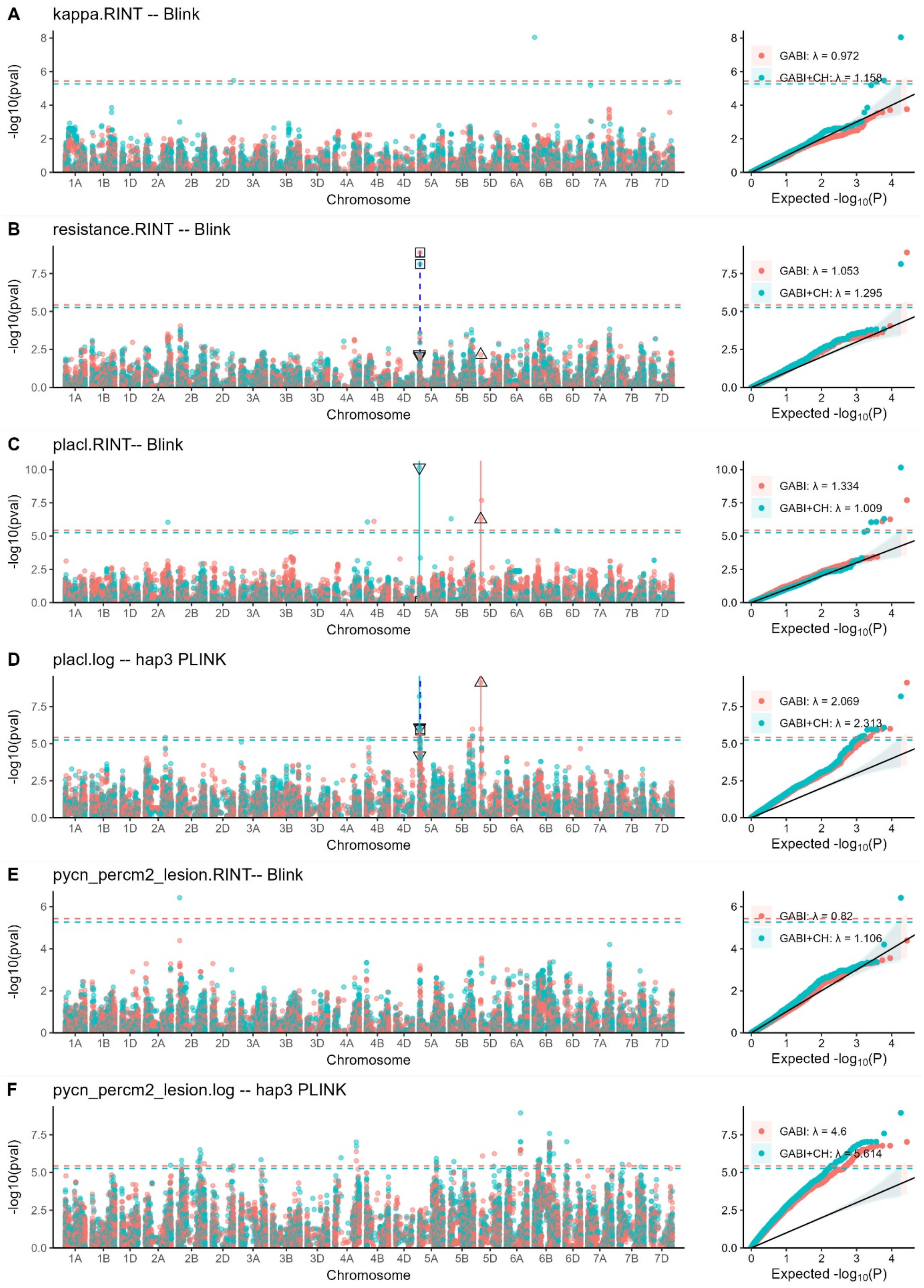


**Supplementary Figure S2: Reproduction of Yates et al. (2019) GWAS outcomes using different models and marker sets.** All analyses were performed using two different sets of genotype data. Red dots represent the original GABI wheat SNP set used in Yates et al 2019, comprising 13,648 SNP for 319 varieties. Blue dots represent the current SNP set including also the Swiss cultivars missing in the GABI set, comprising 9,125 SNP for 330 varieties. Dashed red and blue horizontal lines represent the Bonferroni significance threshold for the respective data sets. Vertical lines together with squares and triangles highlight significant MTAs shared across different datasets. Across all traits, more significant MTA were detected using the current SNP set including the Swiss cultivars. **A-B:** GWAS results (Manhattan and QQ-plots) for the traits kappa and resistance, respectively, from the current study, using rank-based inverse normal transformed (RINT) data and the Blink GWAS model. Note that the MTA for kappa disappear when Swiss cultivars are excluded. The results for resistance are, however, identical. **C:** GWAS results for the trait PLACL (Yates et al., 2019) with RINT transformed data and the Blink GWAS model. **D:** GWAS results for the trait placl (Yates et al., 2019) with log transformed data conducted using the PLINK GWAS model using haplotype blocks comprising three consecutive SNP. Note that the QTL on chromosome 5A overlap between the log (D) and RINT (C) transformed data for the trait PLACL as well as the resistance trait (B). Note also that the PLINK model shows substantial p-value inflation (QQ-plot in (D) compared to the QQ-plot in (C). **E-F:** GWAS results for pycnidia density within lesions (pycn_percm2_lesion, Yates et al., 2019) using RINT-transformed data with the Blink GWAS model log-transformed data with the PLINK haplotype GWAS model, respectively. Note the severe p-value inflation displayed in the PLINK model versus the Blink model (QQ-plot in F versus QQ-plot in E)
